# Timing matters: A meta-analysis on the dynamic effect of stress on salivary immunoglobulin

**DOI:** 10.1101/2024.01.10.575001

**Authors:** Lennart Seizer, Lukasz Stasielowicz, Johanna Löchner

## Abstract

The impact of psychological stress on physiological systems has been a focus of extensive research, particularly in understanding its diverse effects on immune system activity and disease risk. This meta-analysis explores the dynamic effect of acute stress on salivary immunoglobulin-A (S-IgA) levels, a key biomarker for secretory immunity within the oral environment. Analyzing data from 34 samples comprising 87 effect sizes and a total of 1,025 subjects, a multi-level approach is employed to account for the temporal variability in measuring the stress response. The results reveal a significant increase in S-IgA levels peaking around 10 minutes after stress exposure, followed by a return to baseline levels approximately 30 minutes later. In addition, the meta-analysis identified several research gaps of the extant literature, such as limitations in the considered time lag after stress. In conclusion, the findings emphasize the temporal nuances of the stress-induced S-IgA response, which can help to infer potential biological pathways and guide sampling designs in future studies. Further, we highlight the use of a multi-level metaanalysis approach to investigate the temporal dependencies of the interplay between stress and immune functioning.

## 1. Introduction

The increasing interest in the influence of psychological stress on physiological systems has spurred extensive research efforts, improving our understanding of the multifaceted stress effects on on immune system activity and disease risk (Everly et al., 2019). A variety of stressful events has been investigated by previous research, ranging from brief everyday stressors (e.g., academic exams), over life event stressors (e.g., death of loved ones), to standardized laboratory stressors (e.g., Trier social stress test). Across all of these stressors, effects on immune system activity have been found, such as increases in peripheral inflammatory cytokines. These changes may be part of an adaptive fight-or-flight response to stressful situations, which evolutionary carried the risk of injury and infection (Szabo et al., 2020; Marsland et al., 2017; Slavish et al., 2015; Segerstrom and Miller, 2004). Further, psychological stress and the individual degree of stress responsivity are considered risk factors in various somatic and psychiatric diseases, possibly by affecting immune functioning (Seizer and Schubert, 2022; Khandaker et al., 2021; Everly et al., 2019; Marsland et al., 2002). However, the pathways underlying this stress-related regulation of immunity are still not fully understood (Schiller et al., 2021), and the field is plagued by high heterogeneity of effect sizes reported in individual studies (Szabo et al., 2020; Segerstrom and Miller, 2004). Further, current research in psychoneuroimmunology lacks a strong *Theory of Change*, that is for example, a clear understanding of the temporal trajectory of immune responses to stress and to what extent these responses depend on aspects like the effective duration of stress or the temporal lag between stress exposure and measurement of its effects (Moriarity and Slavich, 2023; Seizer et al., 2023; Agorastos and Chrousos, 2022; Rohleder, 2019). Therefore, in the present meta-analysis, we aim to elucidate the role of temporal characteristics in the relastionship between stress and a key component of mucosal immunity – salivary immunoglobulin-A (S-IgA). Since 95% of all infections start at mucosal surfaces such as the mouth and the respiratory tract, and S-IgA concentrations predict susceptibility to such infections, our findings will be of clinical importance as well (Engeland et al., 2016; Nakamura et al., 2006; Tiollier et al., 2005).

Recent meta-analyses have summarized the influence of acute psychological stress on immune system activity: Marsland et al. (2017) investigated the inflammatory reactivity to acute laboratory stress paradigms and found significant stress-related increases in the levels of circulating interleukin (IL)-1*β*, IL-2, IL-6, IL-10, and tumor-necrosis-factor (TNF)-*α*. Notably, none of these increased during the first 10 min following stress. Instead, increases were observed at later times, such as after 40-50 min, with IL-6 showing a prolonged response up to 120 min post-stress exposure. Slavish et al. (2015) and Szabo et al. (2020) both reviewed studies that assessed salivary markers of inflammation following acute laboratory stressors. Slavish et al. (2015) found salivary IL-1*β*, IL-6, and TNF-*α* to increase following stress, but many studies only assessed salivary markers on one or two occasions and only up to 30 min post-stress, which did not allow to further look into the kinetics of inflammatory change in saliva. Similarly, Szabo et al. (2020) found stress-induced increases in IL-6, IL-8, IL-10, TNF-*α*, and IFN-*γ*. Further, they found no moderation by any demographic factors (i.e., age, health status, and sex) of these effects. However, none of these studies investigated the influence of stress on S-IgA, either because no saliva samples were considered or because only markers of inflammation were of interest.

S-IgA is the predominant immunoglobulin found in mucus secretions originating from salivary glands and frequently serves as a biomarker for secretory immunity within the oral environment. Furthermore, it may be representative of IgA-secreting B-lymphocytes across diverse mucosal regions throughout the body (Graham-Engeland et al., 2018; Heaney et al., 2018). S-IgA is synthesized from B-lymphocytes positioned adjacent to mucosal cells, conveyed through intracellular pathways, and subsequently discharged into the secretions of these cells (Brandtzaeg, 2013). This production and release of S-IgA follows a two-step process: Initially, polymeric IgA, as released by B-cells, binds to a receptor molecule known as the polymeric immunoglobulin receptor (pIgR), which is located on the basolateral surface of a secretory glandular cell. Subsequently, the pIgR is transported from the basolateral surface to the apical surface of the glandular cell. Consequently, both the secretion of IgA by plasma cells and the presence of pIgR available for translocation by glandular cells are crucial factors that determine the rate of S-IgA secretion. S-IgA levels are under neuroendocrine control and may be regulated by acute stress via the activation of sympathetic nerves and subsequent release of antibodies from B-lymphocytes (which have a high density of *β*2-adrenergic receptors) and/or an increased transport of IgA across the epithelium into saliva (Engeland et al., 2016; Teeuw et al., 2004; Bosch et al., 2002).

In this meta-analysis, we aim to aggregate the results from studies that have assessed the effect of acute stress on S-IgA. Specifically, we will investigate the standardized mean difference between baseline S-IgA values and values assessed after stress exposure. Crucially, we will examine whether the effect of stress on S-IgA is moderated by the time lag between stressor onset and saliva sampling, i.e., the dynamic time effect. Our multi-level meta-analytic model accounts for the dependency of effect sizes and allows us to include all S-IgA measurements rather than only two times points per sample (Cheung, 2014; Assink et al., 2016). For example, if S-IgA was measured four times after stress exposure, all measurement occasions can be included in the meta-analysis. This approach will allow us to determine a dynamic response trajectory of S-IgA with the timing of onset, peak, and recovery of the effect. This temporal information may help to infer plausible biological mechanisms of stress effects. Further, knowledge about the temporal characteristics can inform the choice of sampling times in study protocols of future research. After all, if sampling time points are inadequate, stress responses may be missed or misjudged (Stoffel et al., 2021; Bolger and Laurenceau, 2013). Beyond laboratory studies, there is a growing interest in the application of intensive longitudinal designs to measure immune activity during everyday situations (Moriarity and Slavich, 2023; Seizer et al., 2022; Schubert et al., 2021), whereby saliva sampling is a promising method for real-life research approaches with ambulatory assessments (Slavish et al., 2015).

## 2. Methods

### 2.1. Inclusion criteria

Only studies meeting the following criteria were included in the meta-analysis: reports on studies that investigated the (1) changes in S-IgA levels in response to (2) acute psychological stress in (3) healthy human subjects and are published in (4) English language. Studies that examined the effect of physical exercise or chronic psychological stress were excluded (e.g., Engeland et al., 2016). The articles found in the literature search were screened and selected for inclusion by two researchers. Consensus for inclusion/exclusion of articles was obtained through discussion.

### 2.2. Literature search

Three databases were searched in May 2023 to find relevant studies: PubMed, PsycINFO, and Web of Science. Stress keywords (e.g., stress and stressor) were combined with S-IgA keywords and abbreviations (e.g., immunoglobulin-A, IgA, saliva, and salivary).^1^ A summary of the literature search is provided in Figure 1. In the database search, 1,789 articles were found. In the first step, the titles and abstracts were screened. When no decision about the relevance of the study could be made, the full text was examined. Thereby, 61 potentially relevant reports were identified for full-text screening. In this second step, 30 more studies were excluded because they did not fit the inclusion criteria (k = 13), were not available (k = 6), were not published in the English language (k = 5), or relevant data could not be obtained (k = 6). Overall, relevant data were identified for 34 independent samples (31 reports).

**Figure 1:**
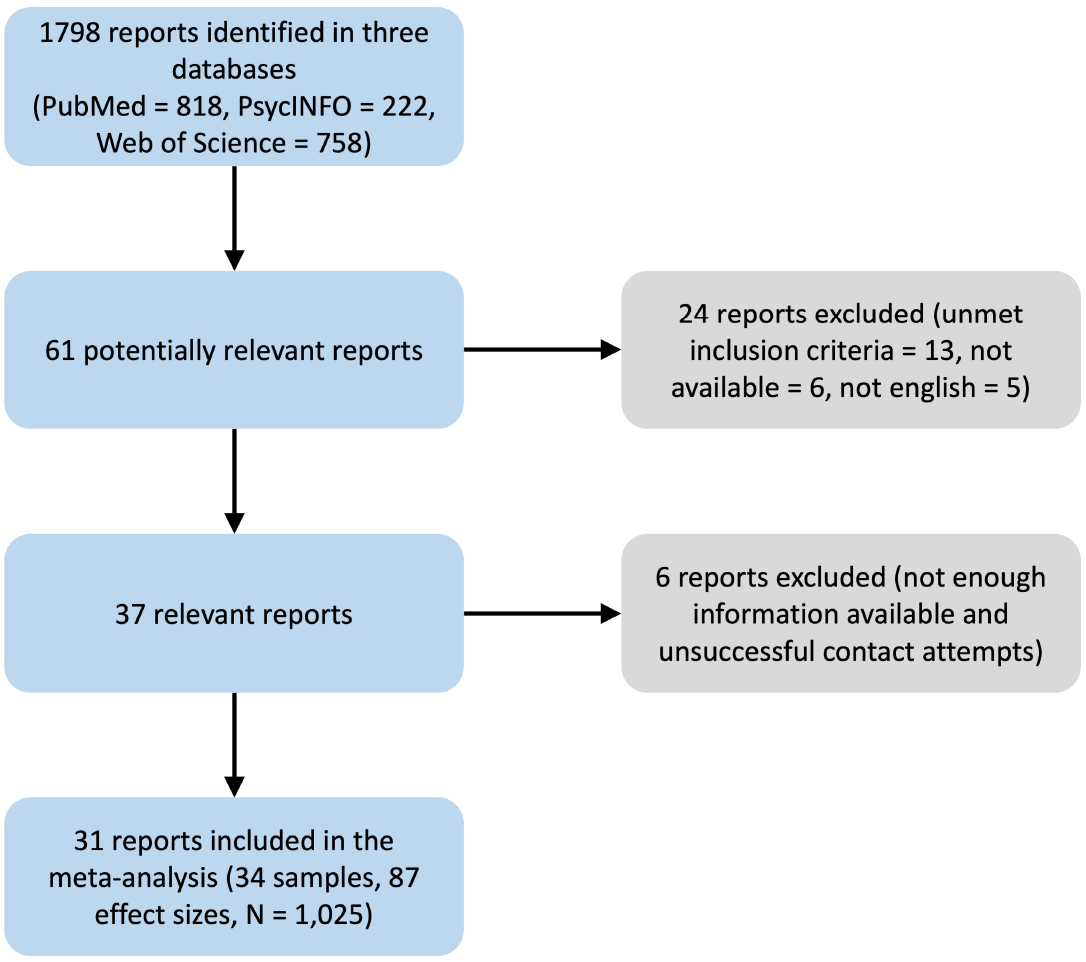
Flow chart summarizing the literature search.

### 2.3. Coding

Each article was coded twice, and any inconsistencies were resolved through discussions among the coders. To determine the effects of stress on S-IgA, the means and standard deviations of S-IgA measurements at different time points were extracted from the articles, along with the information on when exactly S-IgA follow-up measurements were made relative to the baseline measures. The correlations between the measurements at different time points were also coded. If studies investigated the effects of cumulative stress by repeating stress protocols multiple times, only the results from the first experiment were used. Further, possible moderators were coded, such as the method of saliva collection, the normalization procedure for S-IgA concentrations, the type of stressor, the duration of the stressor, the mean sample age, and the proportion of men in the sample. When relevant data could not be obtained from the original article, the corresponding author was contacted. If the corresponding author did not reply and the relevant data was published as a figure, the data was extracted from the figures using the *metaDigitise* package in *R* 4.2 (Pick et al., 2019). If the data necessary to compute effect sizes could not be obtained, the study was not included in the analyses.

### 2.4. Statistical analysis

All analyses were performed in *R* 4.2.3 (R-Core-Team, 2022) using the package *metafor* for meta-analytic modeling (Viechtbauer and Viechtbauer, 2015). Full reproducible code and output for all analyses is available at https://osf.io/fwyzc/. Effect sizes were calculated as the within-group standardized mean difference (SMD) using the mean and standard deviations of S-IgA at baseline and after stress exposure. For the correlation between baseline and followup measures, we assumed a correlation of *r* = .61 because it was the average correlation of the available data. A sensitivity analysis was conducted to assess the robustness of the pooled effects by varying this correlation coefficient (*r* = .4 and *r* = .8), which revealed the same pattern with respect to the statistical significance across the analyses. We used Hedges’ *g* as the effect size in meta-analytic models, as it adjusts effects from small samples to reduce bias. In the present meta-analysis, positive *g* values reflect an increase in S-IgA, relative to the baseline, in response to psychological stress.

Considering that studies measure S-IgA in various intervals, estimating the mean stress response is of limited utility. Therefore, to account for the dynamic nature of stress response, we included time trends in the present meta-analysis. Specifically, for each effect size, the delay between the baseline S-IgA measurement and the S-IgA measurement after stress exposure was coded in minutes. Since the stress response trajectory can be nonlinear and it was unclear which time intervals are used in the extant literature before conducting the literature search, we refrained from arbitrarily assuming a specific trajectory. Instead, we compared 20 natural cubic spline models with increasing complexity, using the Akaike Information Criterion (AIC) to find the most adequate model. Model selection based on information criteria aims to identify the most accurate models while ensuring that they are as parsimonious as possible. After all, adding further variables sometimes leads to only negligible predictive improvements. To identify the best of the 20 candidate spline models in the present meta-analysis, we used the respective AIC values to estimate model weights (wAIC) (Wagenmakers and Farrell, 2004). The model with four degrees of freedom was identified as the best of the 20 considered models (wAIC = .53), followed by a model with five degrees of freedom (wAIC = .22). Therefore, the model with four degrees of freedom was used to model the trajectory of the stress response.

In all meta-analytic models, we used the *t* distribution rather than the less conservative *z* distribution to calculate test statistics. Thereby we reduced the risk of false discoveries. Following the recommendations (Hönekopp and Linden, 2022; Langan et al., 2017), Restricted Maximum Likelihood was used as the estimation method, with the exception of model comparisons, which were based on Maximum Likelihood estimation as the compared models included different predictors (Long and Ryoo, 2010).

#### Dependent effect sizes

In some studies, multiple relevant effect sizes were reported. For example, the stress response was measured on several occasions in the sample, or participants were exposed to various stressors within the study. The way the present meta-analysis approaches dependent effect sizes has substantial advantages over simpler strategies like choosing one effect size per sample or averaging effect sizes within the study (Cheung, 2015; Assink et al., 2016). Specifically, including all relevant effect sizes enables more nuanced data-analytic strategies, such as allowing moderator variables to vary within a study or including all available measurement occasions when estimating meta-analytic effects. Relatedly, including all available effect sizes avoids underestimating the heterogeneity of effect sizes. Furthermore, by avoiding the loss of information, our analyses will have more power.

To account for dependent effect sizes, we used meta-analytic models with a non-diagonal variance-covariance matrix. The variance-covariance matrix is used to weigh the available effect sizes during the estimation of the meta-analytic effect size. Since the lack of dependence between effect sizes based on the same sample is an unrealistic assumption, we allowed the nondiagonal elements of the variance-covariance matrix (i.e., covariances) to be larger than zero when imputing the matrix with the *metafor* package. To account for the fact that effect sizes are more strongly interrelated if the follow-up measures were made within several minutes rather than a few hours, we also included an autocorrelation parameter during the imputation of the matrix. As usual, the diagonal elements of the matrix were based on the sampling variances of the included studies. In general, the larger the sample, the smaller the sampling variance.

In addition to paying attention to the variance-covariance matrix, we took advantage of the multi-level meta-analytic approach (Assink et al., 2016) and included random effects to model data dependency. Including random effects for samples respects the assumption that effect sizes based on the sample size are more similar to each other than effect sizes based on different samples. However, we also added random effects allowing effect sizes to vary within samples. By including these random effects, it is possible to differentiate between true variability of effect sizes and error variability. Specifically, individual effect sizes can deviate from the meta-analytic effect size not only due to sampling error but also due to systematic differences between samples (e.g., different age groups) as well as systematic variability within samples (e.g., use of several stressors).

#### Heterogeneity

To assess the systematic variability between effect sizes in the present metaanalysis, various measures were used. In addition to the visual inspection of the distribution of the effect sizes (see forest plots in the supplementary materials), we assessed the *Q* statistic. A statistically significant value indicates that there is more variability than expected. Furthermore, we inspected the variance of the random effects *τ* ^2^. By taking the square root of *τ* ^2^, it is possible to directly compare the meta-analytic effect sizes to the variability. If *τ* is relatively high, then it means that the meta-analytic effect cannot be generalized to other studies; considerably smaller and larger effects are also possible.

In the current meta-analysis, several potential moderator variables were coded. Most of these variables were categorical: stressor context (laboratory vs. naturalistic setting), stressor type (e.g., academic exam), normalization (e.g., saliva flow rate), saliva collection method (e.g., passive drool), and sample type (e.g., students). Three metric variables were also coded: stressor duration (in minutes), mean sample age (in years), and proportion of men in the sample. To account for the fact that the distance between baseline and follow-up measures varied between effect sizes, we included the time trend variables together with each moderator. Furthermore, interactions between the time trend and each moderator were also included. This enabled us to examine the possibility that trajectories differ between moderator subgroups (e.g., different trends for each stressor type). To facilitate interpretation, we plotted model-based trajectories for each subgroup (see supplementary materials).

#### Publication bias

To examine the possibility that some findings, such as small effects from small studies, are less likely to be published in this research field, we conducted systematic analyses. First, we chose to compare effect sizes reported in journal articles to other sources, such as books and theses. Large differences between the mean effect sizes could indicate publication bias. Second, we conducted a modified Egger’s regression test for asymmetry in the effect size distribution (Cooper et al., 2019; Pustejovsky and Rodgers, 2019). Specifically, we used sample size information 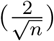 as a moderator. A relationship between this moderator and effect sizes would indicate that effect sizes are distributed asymmetrically, which could be a sign of publication bias (e.g., lack of small effects in small studies).

## 3. Results

### 3.1. Descriptive statistics

In total, 87 effect sizes were available for the 34 samples (*N* = 1,025). Sample sizes tended to be small (*M* = 30.15, *Mdn* = 25, *Min* = 10, *Max* = 92), and most of them were relatively young (*M* = 24.46, *Mdn* = 21.15, *Min* = 19.08, *Max* = 57.40, (*k*_1_ = 22)). The proportion of male participants varied between the samples, but in general, samples consisted of fewer male than female participants (*M* = .36, *Mdn* = .32, *Min* = 0, *Max* = 1). However, it is important to note that sex or gender composition was not reported in seven of the 34 studies (20 %). Included studies are mostly based on student samples (*k*_1_ = 27), but some studies were carried out with other groups, such as children, police officers, or professors. Most studies were conducted in laboratories (*k*_1_ = 25) rather than in a naturalistic context (*k*_2_ = 9). Examples of stressors used in the available studies include academic exams, public speech, and mental arithmetic tasks. Participants were exposed to such stressors for a period between two minutes and two hours (*M* = 21.11, *Mdn* = 14.50). In general, researchers measured S-IgA twice after stressor exposure (*M* = 2.56, *Mdn* = 2.00), but in one study, as many as 10 measurements were made. Typically, the stress response was measured within 30 minutes (*M* = 33.77, *Mdn* = 20). However, various time lags were used (*Min* = 0 minutes, *Max* = 12 hours). In approximately half of the studies, a normalization for S-IgA levels was applied, such as saliva flow rate (*k*_1_ = 18) and total protein (*k*_2_ = 1). Usually, a swab-based approach was used to collect saliva samples (*k*_1_ = 16), but in some samples passive drool (*k*_2_ = 10) or spit samples were used (*k*_3_ = 4).

As expected, S-IgA levels were generally higher after stress exposure than at the baseline (*g* = 0.33, 95% CI [0.15, 0.51], *p* = .001). However, we argue that it is not meaningful to pay particular attention to the magnitude of this average effect size, as it cannot be assumed that the same stress response is shown shortly after stress exposure and multiple hours after exposure. Therefore, we account for the time lag between S-IgA measurements in the following analyses.

### 3.2. Stress response trajectory

As can be seen in Figure 2, the stress response depends on the time of measurement. The largest stress response can be found approximately 10 minutes after exposure; it corresponds to approximately *g* = 0.70. Afterwards, the response decreases rapidly and tends to be undetectable after 30 minutes. Considering this pattern, it is not surprising that adding the four spline terms to model the non-linear trajectory was helpful (*F* (4,76) = 5.06, *p* = .001). However, considerable heterogeneity was found when estimating the average trajectory. Some effect sizes in Figure 2 deviate substantially from the overall trend. There was substantial heterogeneity (*Q* = 22941447.92, *p <*.001) between samples (*τ* = 0.45) and within samples (*τ* = 0.31). Therefore, moderator analyses were conducted to examine possible explanations for this heterogeneity.

**Figure 2:**
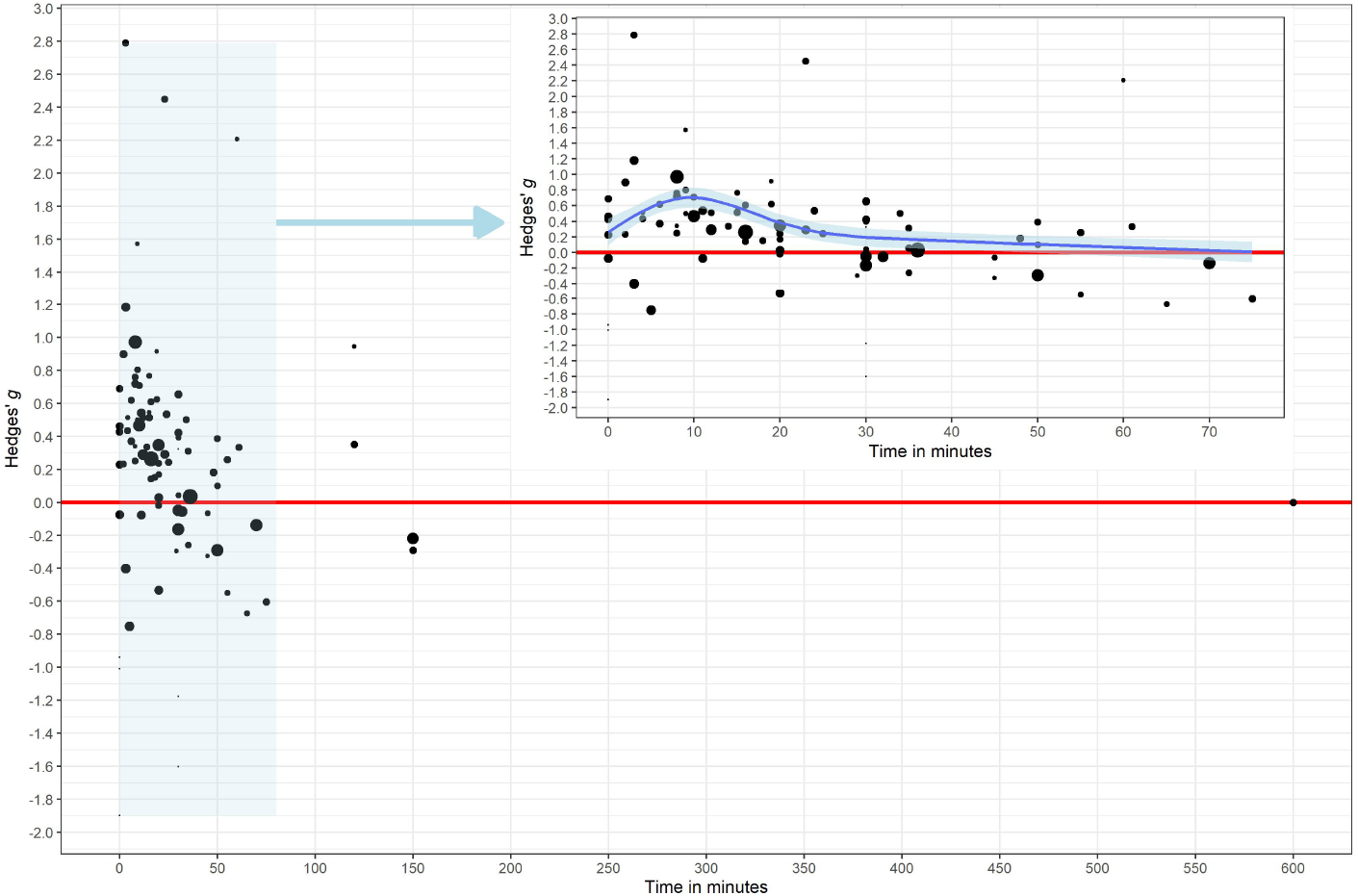
Dependence of the stress response on the measurement time point. The effect size is the standardized mean difference (*g* ) between baseline salivary immunoglobulin-A (S-IgA) and S-IgA measured after exposure to a stressor. In general, the stress response increases in a non-linear fashion in the first minutes, reaching the maximum level after approximately 10 minutes. Stress effects systematically decrease in the following minutes and S-IgA usually returns to the baseline level within one hour. The size of the points reflects the sample size, with larger points corresponding to larger samples. The figure in the upper right corner zooms in on the effect sizes highlighted in blue, which are based on relatively short time lags. To increase the readability, only the zoom perspective includes a trend curve.

### 3.3. Moderator analyses

For several reasons, we restricted the moderator analyses to effect sizes with time lags under 30 minutes. First, as shown in Figure 2, many effect sizes were available within this range (72% of effect sizes), which is necessary to reliably estimate trajectories for different subgroups. Second, the effect sizes within this time range tend to be positive, which is consistent with the expectation that S-IgA increases rather than decreases after being exposed to a stressor. Hence, the effect sizes tend to capture the phenomenon of interest (i.e., S-IgA increase). Third, the average trajectory between 0 and 30 minutes resembles a quadratic trend, which can be modeled using simple quadratic polynomials instead of natural cubic splines to facilitate the interpretation of meta-regression coefficients.

We examined whether the trajectory depends on the stressor context, stressor type, stressor duration, normalization, saliva collection method, sample type, mean sample age, and proportion of men in the sample. No evidence of substantial differences between trajectories was found. However, visual inspection of the trajectories (see https://osf.io/fwyzc/ for all plots) indicated that even the restricted range of time lags was not covered fully in some moderator subgroups. For example, some studies conducted in laboratories did not measure S-IgA immediately after stress exposure, and studies conducted in a more naturalistic context did not use time lags between 10 and 20 minutes after stress exposure. Thus, there is considerable uncertainty about some parts of the trajectories, which precludes making definitive conclusions about the moderators at this stage.

### 3.4. Publication bias

No substantial publication bias evidence was found when using the modified Egger’s regression test (*b* = .28, CI 95% [-1.72, 2.27], *p* = .780. Unfortunately, it was not possible to conduct the second planned analysis and approximate publication bias by comparing effects reported in journal articles with other sources because all included studies were journal articles.

## 4. Discussion

This meta-analysis included 34 samples with 87 effect sizes and a total of 1,025 people to investigate the dynamic effect of acute stress on S-IgA levels. Generally, we found the levels of S-IgA to increase following the exposure to acute stress (*g* = 0.33, 95% CI [0.15, 0.51]). However, we recommend refraining from emphasizing the magnitude of the average effect. This advice follows from the assumption of a dynamic stress response, where effects observed shortly after stress exposure may differ from those recorded several hours later. Our findings are also consistent with this assumption. Specifically, after accounting for the time trend, we found the following pattern: acute stress was found to be followed by significant increases in S-IgA with a peak after about 10 min (approximately *g* = 0.70) and a return to baseline after about 30 min. To put this finding into perspective, S-IgA increases observed in the present metaanalysis lie temporally in-between increases in salivary alpha-amylase that reflect sympathetic nervous system activity and are observed within a few minutes after stress exposure (Nater and Rohleder, 2009) and increases in cortisol levels that reflect hypothalamic–pituitary–adrenal (HPA) axis activity and are observed after 15-20 minutes (Russell and Lightman, 2019).

It is important to note that the identified time trend reflects an average trajectory. We conducted moderator analyses to offer a more nuanced perspective and explain the remaining effect size heterogeneity. However, we found no evidence of moderating influences on the stress-induced S-IgA response trajectory by stressor context, stressor type, normalization procedure, saliva collection method, sample type, mean sample age, and proportion of men in the sample. As noted in the introduction, the increase in S-IgA levels in response to acute stress may be attributed to two pathways which can both be enhanced by autonomic nervous system transmitters, that is (1) the redistribution into saliva via increased transport of IgA across the epithelium and (2) the increased production and release of IgA from B-lymphocytes (Engeland et al., 2016; Bosch et al., 2002). Our findings indicate that the stress-induced changes in S-IgA happen quite fast, with a peak after 10 min. This response is likely attributed to the discharge of antibodies that have already been synthesized by plasma cells and enhanced transport of antibodies through the epithelial barrier and into saliva (redistribution effect), given that the temporal delay of the effects does not allow for the production of a substantial quantity of new antibodies (Segerstrom and Miller, 2004; Bosch et al., 2002). Fittingly, studies in rats have demonstrated that stimulating the local autonomic branches results in a rapid increase of S-IgA into saliva, occurring within minutes (Matsuo et al., 2000; Proctor et al., 2000; Carpenter et al., 1998). Combining our meta-analytic S-IgA response trajectory (Figure 2) with these previous reports, the findings suggest that this stress-related autonomic stimulation primarily affects the IgA translocation process, with little impact on IgA release by B-lymphocytes (Teeuw et al., 2004; Proctor and Carpenter, 2002; Carpenter et al., 1998).

To our knowledge, this is the first meta-analysis on the effects of stress on immune system activity to account for the continuous nature of time; previous analyses averaged or binned effect sizes from multiple time points into time categories. The multi-level approach employed in this study provides a robust framework for investigating dynamic effects, offering a significant advantage over previous approaches as it improves the understanding of the temporal nuances of stress’s influence on the immune system. Further, some research gaps and limitations were identified in the included studies, such as (1) the absence of more specific data regarding IgA subtypes and secretory components that would allow further insights into potential mechanisms (Engeland et al., 2016), (2) underrepresented populations, such as children and old people, and relatively small sample sizes of the included studies, (3) only a few studies considered time lags above 30 min, limiting estimation accuracy of the IgA response trajectory at higher lags, (4) different measurement time points were applied between studies, which complicates explaining the heterogeneity of effect sizes between studies, and (5) many studies did not normalize S-IgA levels although it is currently considered preferential practice (Engeland et al., 2019). Future primary studies should aim to address these limitations by obtaining or analyzing more specific data to enhance the understanding of the mechanisms behind the findings of the present metaanalysis.

Conclusively, we provided meta-analytic evidence for stress-induced increases in S-IgA levels. Crucially, accounting for the temporal distance between baseline and post-exposure measures allowed us to estimate the dynamic trajectory of this response. These findings improve our understanding of the interplay between acute stress and immune functioning, emphasizing the importance of temporal considerations in future studies and clinical applications. The determined peak and recovery times of stress responses will help guide researchers to set up the timing of post-stress measurements in future laboratory and naturalistic studies interested in S-IgA responses. Building on the estimated response trajectory, sampling protocols can be created to time the measurements of stress responses optimally. Thus, this meta-analysis not only elucidates the dynamic response trajectory of salivary immunoglobulin-A to acute stress but also provides valuable insights for optimizing the timing of post-stress measurements in laboratory and real-life research settings.

## Footnotes

1 (stress* OR distress) AND (immunoglobulin* OR ig OR sig OR s-ig) AND (saliva* OR oral)

